# InSituCor: a toolkit for discovering non-trivial spatial correlations in spatial transcriptomics

**DOI:** 10.1101/2023.09.19.558514

**Authors:** Patrick Danaher, Dan McGuire, Michael Patrick, David Kroeppler, Haiyan Zhai, Joachim Schmid, Joseph M. Beechem

## Abstract

Spatial transcriptomics presents the best kind of problem: how to find the many biological insights hidden within complex datasets. Spatially correlated genes can reveal high-interest phenomena like cell-cell interactions and latent variables. We introduce InSituCor, a toolkit for discovering modules of spatially correlated genes. A major contribution is that InSituCor returns only correlations not explainable by obvious factors like the cell type landscape; this spares precious analyst effort for non-trivial findings. InSituCor supports both unbiased discovery of whole-dataset correlations and knowledge-driven exploration of genes of interest. As a special case, it evaluates ligand-receptor pairs for spatial co-regulation.

## Background

Single cell spatial transcriptomics data, in which hundreds to thousands of genes are measured in situ across potentially millions of cells, poses a daunting version of the central challenge of ‘omics: how can an analyst possibly discover all the interesting biology contained within a dataset? One class of exploratory analyses looks for spatially correlated sets of genes; that is, genes that tend to be expressed in the same regions^1,2^. Spatial correlation between genes can arise through direct cell-cell communication, or from some underlying latent variable; both these mechanisms are of interest.

Spatial correlation has a limitation: most cell types are spatially organized, which induces spatial correlation among genes whose expression varies across cell types. Thus spatial correlation often provides little more than an oblique readout of the cell type landscape. This pitfall can be avoided by looking one cell type at a time, but this solution misses interactions among multiple cell types. Some methods^3^ wrap cell type into highly specific models; this approach, however, limits the diversity of trends they can discover.

Here we introduce InSituCor, a toolkit for quickly identifying spatial correlations deserving scarce analyst attention. InSituCor identifies gene modules with spatial correlations that cannot be explained by trivial factors like the cell type landscape or technical effects.

## Results and discussion

We demonstrate InSituCor in a colon tumor profiled with a 6000-plex CosMx^4^ panel (Fig 1a). InSituCor begins by taking the expression profile of a neighborhood around each cell (Fig 1b), building an “environment expression matrix” (Fig 1c). Typical methods produce results akin to taking the correlation matrix of the environment expression matrix (Fig 1d). To eliminate the influence of unwanted variables like cell type, signal strength and background intensity, InSituCor builds an “environment confounder matrix” (Fig 1e) summarizing these variables for each cell’s neighborhood. InSituCor defines spatial correlation as the correlation matrix of the environment expression matrix, conditional on the confounder matrix (Fig 1f), or

**Figure 1:**
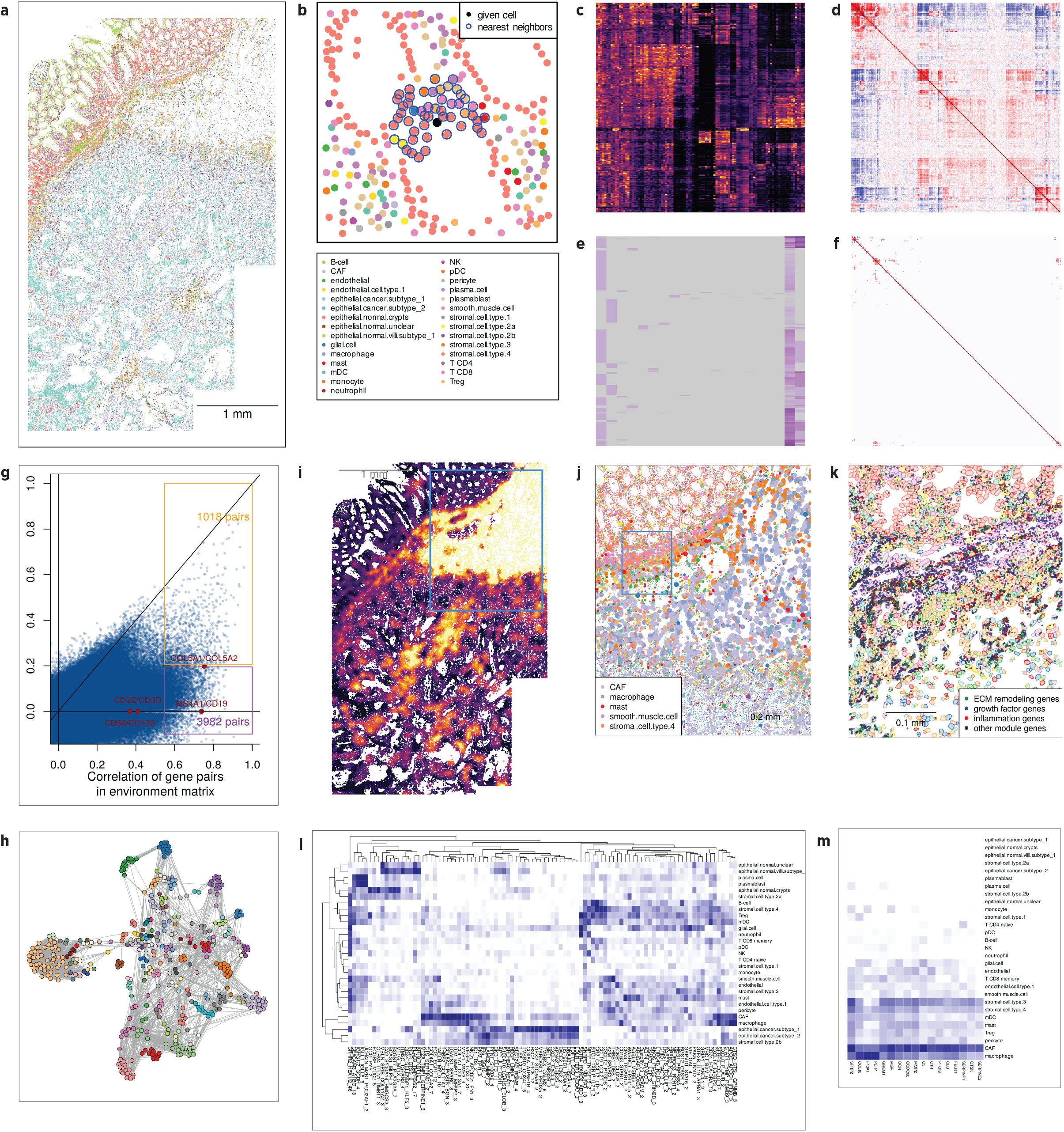
Demonstration of InSituCor workflow. **a**. Cell type map of a colon cancer. Color legend applies to panels a,b,j. **b**. Example of a cell’s nearest neighbors, used to define its “environment expression” profile. **c**. Subset of the environment expression matrix. **d**. Raw correlation matrix of the environment expression matrix showing near-ubiquitous correlations (subset of 300 random genes). **e**. Subset of the environment confounding matrix, encoding cell type abundance and other confounding variables in each cell’s neighborhood. **f**. Correlation matrix of the environment expression matrix (c) conditional on the confounding matrix (e), over the same subset of genes as (d). Most unconditional correlations (d) are fully explained by the confounding variables. **g**. Raw vs. conditional correlation of environment gene expression. Selected pairs of marker genes are highlighted. **h**. Network representation of correlation between all genes in all modules. Genes with correlation > 0.2 are connected. The module from i-k is highlighted. **i**. Environment scores for a “tumor-promoting inflammation” module. **j**. Single-cell scores for the module; point size reflects cells’ module score. The window highlighted in (i) is shown. **k**. mRNA molecules of module genes. The window highlighted in (j) is shown. **l**. Estimated involvement of each cell type in each module. **m**. Estimated involvement of each cell type in each gene of the highlighted module.

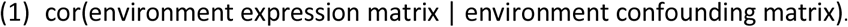

This conditional correlation matrix is the cornerstone of InSituCor analyses. It measures genes’ tendency to be expressed in the same neighborhoods, beyond what cell type and other confounders can explain. Due to the large numbers of cells analyzed, even small correlations attain strong statistical significance; for this reason, InSituCor does not compute p-values, as statistical significance here is a low-specificity indicator of interesting results.

The conditional correlation approach finds that most of the strongest spatial correlations in the unadjusted analysis are spurious: among the top 5000 gene pairs found without adjusting for confounders, with a range of (0.62, 0.97), only 1018 had correlations > 0.2 in the conditional correlation matrix (Fig 1g). Pairs of marker genes, for example CD3D/CD3E and CD19/MS4A1, had strong spatial correlations but near-zero conditional correlations.

InSituCor aids comprehension by extracting modules of mutually co-expressed genes (Fig 1h). One module discovered in this analysis consisted of 17 genes collectively suggestive of tumor-promoting inflammation (Fig 1h,i,j,k,m). This module included genes involved in microenvironment remodeling (CCL18, MMP2, CSTK), growth factor signaling (SFRP2, GREM1, DCN, SERPINF1), and inflammation (C3, C1R, PTGIS).

Each module is scored with a weighted average of its genes; scores are calculated both for cells’ environments and for single cell expression. A map of scores for the tumor-promoting inflammation module shows it peaking in the stroma, with smaller hotspots in the tumor bed (Fig 1i). Looking at single-cell scores for the module, we see CAFs and macrophages driving module activity, with nearby mast cells, smooth muscle cells and stromal cells also participating (Fig1j). Zooming in to resolve individual mRNA’s, we see more nuanced behavior of the module genes across cell types and space (Fig 1k).

Dozens of modules will be discovered in a typical study of ≥1000 genes. To help analysts prioritize, InSituCor estimates the role of each cell type in each module. Cell type involvement is summarized at the module (Fig 1l) and at the gene level (Fig 1m).

In a typical study, the analyst will choose confounding variables, then derive modules with a single R command. Summary plots (Fig 1l,m) will suggest a few modules of particular interest. The analyst can then invest real effort, carefully examining spatial plots (Fig 1k) to develop a nuanced understanding of modules’ behavior.

To speed computation time, many of InSituCor’s calculations use subsets of 5,000 cells. InSituCor took 2.5 minutes on a r5.12xlarge EC2 instance to analyze this dataset of 112,846 cells and 6,000 genes.

InSituCor also supports a knowledge-driven workflow: one simply examines the conditional correlation structure around genes of prior interest. To describe this tumor’s signaling environment, we re-analyzed just the dataset’s 407 ligands (cite cellChatDB). This produced 18 modules containing 51 ligands, many arising from multiple cell types (Figure 2a,b). Focusing on modules involved in the anti-tumor immune response, we see a module of the chemoattractants CCL19 and CCL21 concentrated in a narrow band at the tumor periphery, a module of MHC2 antigen presentation genes diffusing slightly beyond this band, and a module of MHC1 antigen presentation genes peaking in the same region but extending further yet into the tumor bed (Fig 2c-e). This suggests an interpretation in which a core of chemoattractant expression attracts antigen presenting cells, and an adaptive immune response radiates from this core, eliciting MHC1 expression from surrounding cells.

**Figure 2:**
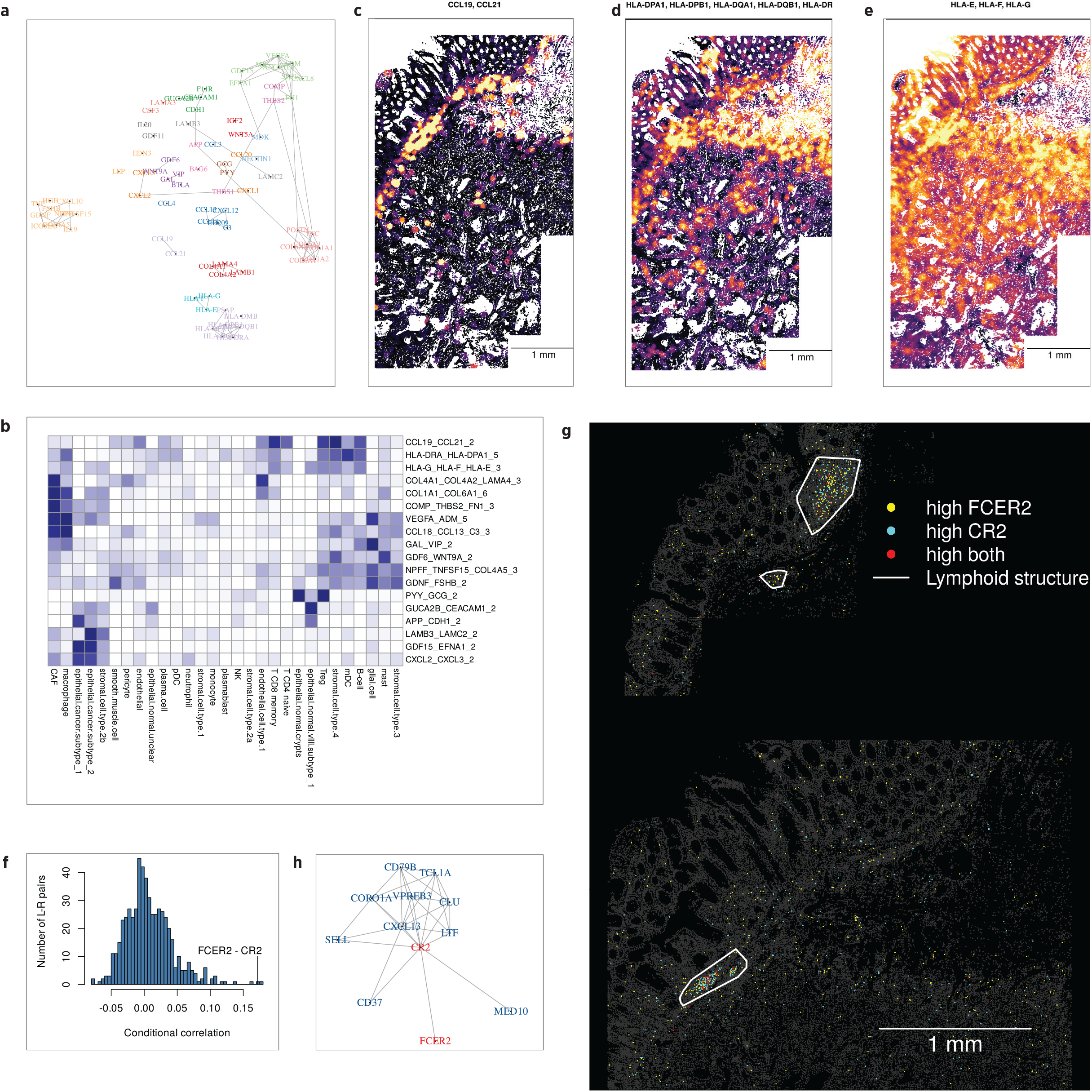
Biology-first use of InSituCor. **a**. Correlation structure of 51 ligands assigned to modules. Edges show conditional correlations > 0.1, and color shows module membership. **b**. Involvement of each cell type in each module. **c**,**d**,**e**. Environment expression of a modules holding chemoattractants (c), MHC2 antigen presentation genes (d), and MHC1 antigen presentation genes (e). **f**. Conditional correlations of 555 ligand-receptor pairs. **g**. Spatial map of single-cell expression of the ligand-receptor pair FCER2 & CR2. **h**. Conditional correlation network around the FCER2-CR2 ligand-receptor pair.

Ligand-receptor pairs motivate another use case, as spatial correlation suggests co-regulation, presumably by the ligand increasing the receptor’s expression or via some latent variable inducing regional expression in both genes^2^. Of the 555 ligand-receptor pairs in this panel^5^, only 11 had conditional correlation > 0.1 (Fig 2f). One correlated pair was FCER2 and CR2, both primarily expressed by B-cells. Visual examination showed their spatial variability to be driven by lymphoid structures (Fig 2g), where B-cells had 2.57-fold (95% confidence interval 1.63-3.51) higher FCER2 and 3.43-fold (2.12-4.75) higher CR2 than B-cells elsewhere.

InSituCor can also be used to explore individual genes of high prior interest. Motivated by the above results, we examined the correlation network around FCER2 and CR2 (Fig 2h). FCER2 had no further connections, but CR2 belonged to a densely-connected network of 10 additional genes involved in B-cell development, activation and trafficking^6-8^. This suggests the hypothesis that the genes connected to CR2 are activated downstream of FCER2 – CR2 signaling.

## Conclusions

Cell type landscapes induce strong spatial correlation between genes, even when those genes do not vary within a cell type. In our example dataset, most of the strongest spatial correlations in the unadjusted analysis proved to be uninteresting after adjusting for cell type abundance. By conditioning on cell type and other confounders, InSituCor ignores these trivial trends and instead reports only correlations indicating more interesting biology. It quickly – in both computational time and analyst time – isolates and summarizes spatial correlations deserving further investigation.

The InSituCor R package is available on request and will be made available at https://github.com/Nanostring-Biostats/InSituCor.

## Methods

Environment expression and confounder matrices are defined by averaging gene expression (or confounder variable values) across each cell’s neighbors. The covariance of the former matrix conditional on the latter is calculated with:

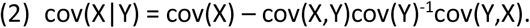

This is equivalent to regressing the environment expression matrix on the confounders matrix and taking the covariance of the residuals. Conditional correlation is calculated by rescaling this covariance matrix to have unit diagonal. The above formula holds for multivariate normal variables; because our environment expression matrix is produced by averaging each cell’s 50 nearest neighbors, the central limit theorem provides some assurance that multivariate normality approximately holds.

To define modules, InSituCor creates a network graph connecting genes with conditional correlations above a threshold, then clusters this graph using the Leiden algorithm^9^. Module scores are calculated as weighted averages of their genes; the default uses inverse square root weighting to account for the Poisson-like mean-variance relation seen in count data.

To calculate “cell type attribution scores”, we first measure the role each cell type plays in each module gene. We summarize this with the correlation between two quantities: first, cells’ environment scores for the module, and second, the cells’ neighborhood expression of the gene in question by the cell type in question. We then summarize the role of the cell type within the module by taking its maximum value of the above statistic across module genes. This produces high scores for cell types that contribute heavily to any of a module’s genes.

A 13mm x 12mm x 5um section of colorectal adenocarcinoma was profiled using the standard CosMx RNA protocol and a 6000-plex, pre-commercial version of the CosMx 6K Discovery RNA panel. 73 fields of view were placed according to markup of a serial hematoxylin and eosin (H&E) stain, focusing on normal intestinal mucosa, lymphoid aggregates, and cancer. The slide was imaged with a 5-channel morphology panel (PanCK, CD68, CD298/B2M, CD45, DAPI). Cell types were defined using the Insitutype R package^10^.

Ligands and ligand-receptor pairs were taken from CellChatDB^5^. Tertiary lymphoid structures were defined by clustering B-cell locations with dbscan^11^. The 3 largest clusters (130, 501 and 905 B-cells) were called tertiary lymphoid structures; the next-largest cluster had just 28 B-cells.

